# A horizontally transferred autonomous Helitron became a full polydnavirus segment in *Cotesia vestalis*

**DOI:** 10.1101/132399

**Authors:** Pedro Heringer, Guilherme B. Dias, Gustavo C. S. Kuhn

**Author notes:** **Corresponding author:** Gustavo Campos e Silva Kuhn, Laboratório de Citogenômica Evolutiva (L3 159), Departamento de Biologia Geral. Instituto de Ciências Biológicas, Universidade Federal de Minas Gerais. Av. Antônio Carlos 6627, CEP: 31270-901, Belo Horizonte, MG, Brazil.

## Abstract

Bracoviruses associate symbiotically with thousands of parasitoid wasp species in the family Braconidae, working as virulence gene vectors, and allowing the development of wasp larvae within hosts. These viruses are composed by multiple DNA circles that are packaged into infective particles and injected together with wasp's eggs during parasitization. One of the viral segments of *Cotesia vestalis* bracovirus contains a gene that has been previously described as a helicase of unknown origin. Here we demonstrate that this gene is a Rep/Helicase from an intact Helitron transposable element that covers the viral segment almost entirely. We also provide evidence that this element underwent at least two horizontal transfers, which appear to have occurred consecutively: first from a Drosophila host ancestor to the genome of the parasitoid wasp *Cotesia vestalis* and its bracovirus, and then from *C. vestalis* to a lepidopteran host (*Bombyx mori*). Our results reinforce the idea of parasitoid wasps as frequent agents of horizontal transfers in eukaryotes. Additionally, this Helitron-bracovirus segment is the first example of a transposable element that effectively became a whole viral circle.

## INTRODUCTION

The family Polydnaviridae is composed of symbiotic viruses exclusively associated with more than 40,000 parasitoid wasp species from two families: Ichneumonidae and Braconidae (superfamily Ichneumonoidea). Polydnaviruses (PDVs) exist both as proviral copies in the wasp genome and as their functional form composed of multiple dsDNA circles packaged into infective particles (reviewed in Gundersen-Rindal *et al.* 2013). During wasp oviposition, PDVs are injected into the host, where they express virulence genes that alter insect physiology and allow the development of parasitoid larvae (Strand and Burke 2013, Drezen *et al.* 2014). PDVs are classified in two genera, Ichnovirus (IV) and Bracovirus (BV), which are in turn associated with ichneumonid and braconid wasps, respectively. Despite similarities between these two groups of viruses, they originated from independent viral integration events on each of these two wasp lineages (Herniou *et al.* 2013).

There are more than 18,000 braconid species described to date, but some estimates point to a total number exceeding 40,000 (Quicke 2015). Braconid wasps are parasitoids of a wide range of insect orders, and are one of the most successful insects used in biological control programs (Wharton 1993). An important species from this family is *Cotesia vestalis*, who has been found parasitizing Lepidoptera species in various regions of Asia, Europe, Africa and the Americas (Furlong *et al.* 2013). Also, *C. vestalis* is one of the main parasitoids of the diamondback moth *Plutella xylostella*, which is a pest of *Brassica* plants that costs annually US$4-5 billion worldwide in crop loss and management (Zalucki *et al.* 2012). For those reasons, *C. vestalis* has been successfully used in numerous biological control introductions (Furlong *et al.* 2013).

The complete genome sequencing of *Cotesia vestalis* bracovirus (CvBV) revealed 157 ORFs distributed in 35 encapsidated segments, with most of these genes also having homologs in other BVs (Chen *et al.* 2011). In the same work, the authors noted that a segment from CvBV (CvBV_c35) encodes a protein displaying similarity with the human Pif1 helicase, which has no sequence homology with any other gene described in BVs. The Pif1 family is a member of the superfamily 1B helicases and is involved in many replication related functions in eukaryotes (Bochman *et al.* 2010). Because all PDVs analyzed to date only express replication genes in the calyx cells of wasps, and do not replicate in host cells (Gundersen-Rindal *et al.* 2013, Bézier *et al.* 2009), the function of this helicase in a CvBV encapsidated segment is worth of investigation, as noted by Chen et al. (Chen *et al.* 2011).

In the present study, we found that segment CvBV_c35 is almost entirely made by a rolling-circle Helitron transposon, which has fused to the CvBV genome. Although there has been reports of transposable elements (TE) (including Helitrons) in BV genomes (e.g. Drezen *et al.* 2006, Thomas *et al.* 2010, Dupuy *et al.* 2011, Guo *et al.* 2014, Coates 2015), we describe for the first time a case where an autonomous TE effectively became a viral segment. Based on our results we propose a scenario to explain the presence of this Helitron in the virus, and suggest the occurrence of two consecutive horizontal transfer (HT) events involving this TE across species from three insect orders: Diptera (oriental Drosophila), Hymenoptera (*C. vestalis*), and Lepidoptera (*Bombyx mori*). Finally, we discuss the conservation of this Helitron as a PDV segment, the possibility of a co-option, and the implications of our findings for the study of HT events.

## MATERIALS AND METHODS

### Rep/Hel amino acid analysis

To confirm that the CvBV_c35 codes a Rep/Hel we first analyzed its amino acid sequence (AEE09607.1) using the NCBI's conserved domain database (CDD) (Marchler-Bauer *et al.* 2015). Rep/Hel ORF sequences were retrieved from Repbase (Jurka *et al.* 2005), either directly or using the CENSOR tool (Kohany *et al.* 2006) with CvBV_c35 ORF as a query. Sequences were then aligned using the MUSCLE alignment on MEGA7 (Kumar et al. 2016). We determined the residues that characterize Rep domains by visually inspecting the CvBV_c35 amino acid sequence, based on the consensus from Kapitonov and Jurka (Kapitonov and Jurka 2007) (Figure 1C).

**Figure 1.**
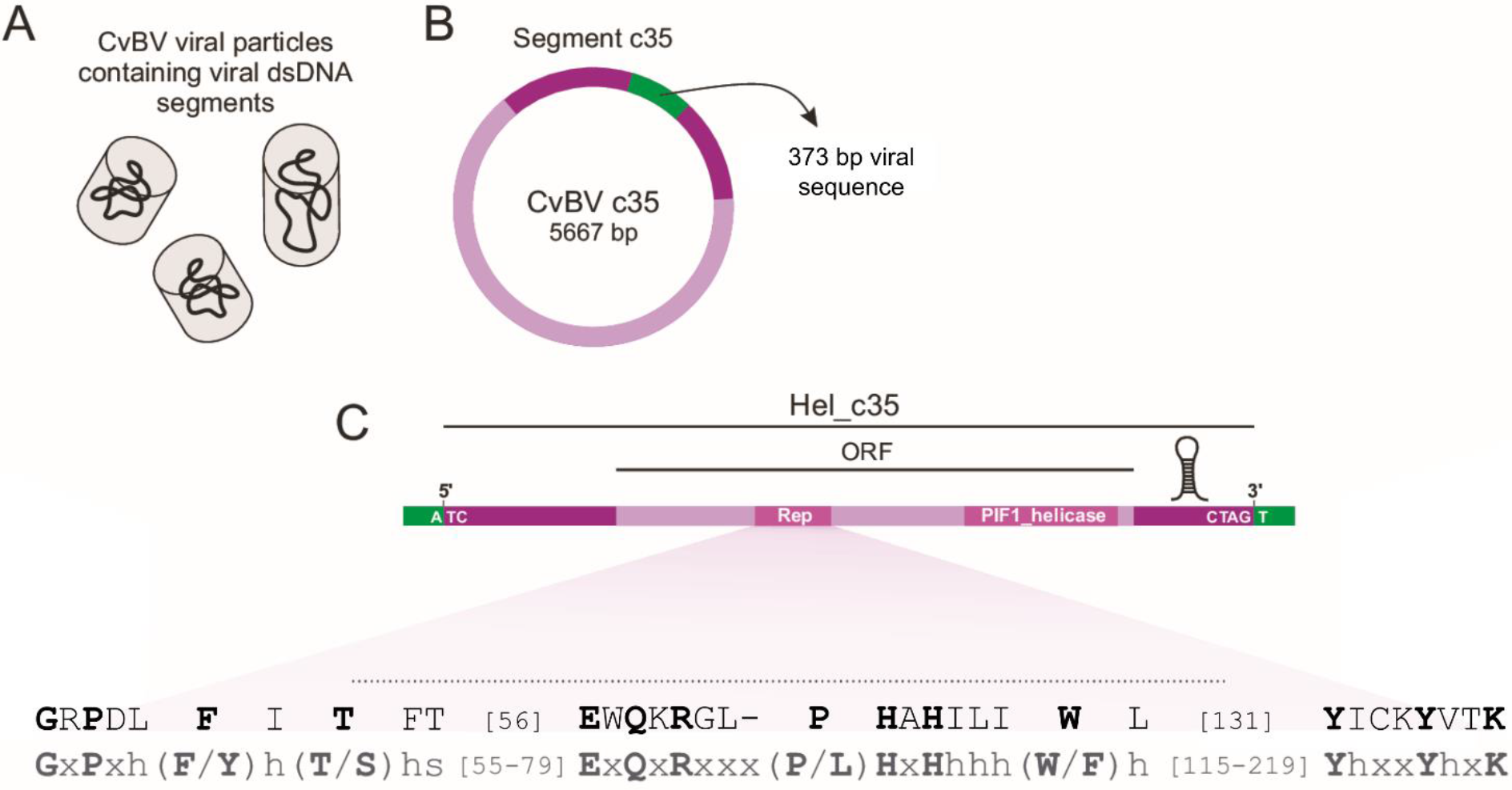
Schematic diagram of the Helitron-containing segment c35 from CvBV. (A) CvBV segments are encapsidated as double-strand DNA circles. (B) Segment c35 contains an ORF (light purple) flanked by sequences similar to Helitron TEs (dark purple). This segment also has a 373 bp viral sequence (green). (C) Structural and coding features of Hel_c35. The Rep catalytic core residues are depicted in black, and the Helitron consensus Rep domain (from Kapitonov and Jurka 2007) is shown below in grey. Conserved residues in the consensus sequence are shown as capital letters. Hydrophobic, small and variable residues are represented by h, s and x, respectively.

### Hel_c35 structure hallmarks

To determine the whole Helitron structure we first used the CvBV_c35 sequence as a query in a search against the Repbase reference collection (Kohany *et al.* 2006). After visually establishing the element's putative boundaries (Figure S1), we determined its precise termini by aligning CvBV_c35 with the best results, and verified if the sequence contained the hallmarks of Helitrons, like the insertion between AT nucleotides, the hairpin structure close to the 3'-end, and the conserved terminal nucleotides. To further validate the TE limits, we used the putative Helitron sequence (named Hel_c35) as a query for a Blastn search (Altschul *et al.* 1990) against the genome of *Cotesia vestalis* available on GenBank (Benson *et al.* 2013), and then used the best results, together with their flanking sequences as a query to a second Blastn search. Both searches were conducted using the following default parameters: Expect threshold 10; Word size 11; Match/Mismatch Scores 2, −3; Gap Costs Existence 5, Extension 2. Most results with multiple hits only display similarity with Hel_c35 sequences within the terminal nucleotides, suggesting that our defined limits for the element encompass the whole TE.

### Search for CvBV_c35 proviral locus and analysis of PDV conserved sequences

To Blast-search (Altschul *et al.* 1990) the CvBV_c35 proviral locus, we used the sequence of this segment (HQ009558.1) as a query against the *C. vestalis* genome on GenBank (Benson *et al.* 2013) using default parameters and selecting the best result based on score, query cover and identity. We then scanned the selected contig (gb: JZSA01007369.1) searching for sequences displaying similarity with PDV genomes and other PDV-related sequences available on Genbank (Benson *et al.* 2013). To detect structural features on the PDV-like sequences found in the selected contig, we first used dotplots from the software Dotlet (Junier and Pagni 2000), followed by manual curation of sequences. Finally, the analysis of palindromic or inverted repeat motifs, and their predicted secondary structures, were made using the softwares mfold (Zuker 2003) and Palindrome analyser (Brázda *et al.* 2016).

### Transcriptome analysis of the Asiatic rice stem borer, *Chilo suppressalis*

To further investigate the presence of the Hel_c35 in *Cotesia* bracoviruses apart from the ones with available genomes, we searched for any sequence data generated from transcriptome studies. The only work we found which was suitable for this analysis was conducted by Wu et al. (2013) on the lepidopteran host *Chilo suppressalis* (Asiatic rice stem borer) while parasitized by *Cotesia chilonis* (see *Results and Discussion*). We used the Rep/Hel coding DNA sequence (CDS) of CvBV_c35 (HQ009558.1) as a query for Blastn searches against the Short Read Archive (SRA) (SRR651040) and Transcriptome Shotgun Assembly (TSA) (GAJS00000000) generated by the same study (Wu et al. 2013). Reads with identity > 90%, and Unigenes with score ≥ 200 were selected. The sequences of selected Unigenes were then used as queries for Blastn searches against genomes from all insect orders for comparison. All searches were conducted using default parameters. The sequences from Unigene42046 and the corresponding region from the CvBV_c35 CDS were aligned using M-coffee (Moretti *et al.* 2007) (Figure S3).

### Search on Arthropoda genomes, alignment and phylogeny

We conducted Blastn searches (Altschul *et al.* 1990) on all Arthropoda genomes available on GenBank (Benson *et al.* 2013), using a query of 1,675 bp from the Hel_c35 Rep/Hel ORF located in the ‘Rep region’ and applying default parameters (see topic ‘*Hel_c35 structure hallmarks*’ above). This region contains the Rep domain that is exclusive from Helitrons (File S1), in contrast to the ‘Hel region’ and its Hel domain belonging the Pif1 family of helicases, which are involved in many cellular processes and pervasive in eukaryotes (Bochman *et al.* 2010). Therefore, to avoid false positives, and because most of the initial sampling searches using the whole ORF (4,538bp) retrieved only fragmented hits, we decided to use a shorter query covering only the Rep region. This selected nucleotide stretch corresponds to the “Helitron_like_N” domain annotated on the NCBI's CDD (Marchler-Bauer *et al.* 2015), the rest of the catalytic core that we manually selected using the Helitron consensus (see ‘*Rep/Hel amino acid analysis*’ above), and an intermediate region between Rep and Hel (File S1). In order to include in our study only TEs from the same family (see Wicker *et al.* 2007 for a hierarchical proposal for transposable elements classification), results with > 70% cover and > 80% identity were selected based on their Max score, and then aligned using MUSCLE on MEGA7 (Kumar *et al.* 2016). The final alignment with ~ 838 bp was formatted on Jalview (Waterhouse *et al.* 2009) to be better visualized (Figure S4). Although we recognize that Helitrons families should be classified in a manner distinct from most DNA TEs, due to their peculiar structure and evolutionary dynamics (Thomas and Pritham 2015), our interest was to retrieve coding sequences, including the ones from elements lacking their terminal 30 bp used in the mentioned classification. To build the phylogeny of aligned Rep sequences, we used the Bayesian inference method on MrBayes 3.2 (Ronquist *et al.* 2012) running a mixed analysis, which samples across the different nucleotide substitution models. The results were compared with the phylogeny estimated using the Maximum Likelihood method on MEGA7 (Kumar et al. 2016) and applying the Tamura 3-parameter model (Tamura 1992), which resulted a similar branch topology (data not shown). The main geographical distributions of the taxa included in the phylogeny were retrieved from various Web sources.

### Date estimation of HT events

For the date estimation of the HT events between insect species, we used the equation for divergence time given by

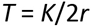

where *T* is the number of generations, *K* is the number of substitutions per site, and *r* is the rate of nucleotide substitution, which is equal to the mutation rate (*μ*) for neutral mutations (Graur and Li 2000). The sequences used comprised the 5’ and 3’ non-coding regions, which flank the Hel_c35 ORF and together sum ~ 750 bp. The copy from CvBV was used to represent *C. vestalis*, and as a query for Blast searches in the genomes of *D. rhopaloa* and *B. mori* (see *Results and Discussion*). Although we used non-coding sequences flanking the Hel_c35 ORF for the analysis, assuming they evolve neutrally, the importance of these regions for Helitron transposition are not fully understood, apart from the terminal 40 bp on each end (Grabundzija *et al.* 2016). As the selective pressure on these sequences cannot be discarded, the assumption of neutrality should be taken cautiously. For that reason, we used one equation (*T* = *K*/2*μ*) for a minimum date estimation, assuming both sequences evolved neutrally, and another (*T* = *K/μ*) for a maximum date estimation, assuming only one sequence evolved neutrally. This second equation (maximum threshold) gives the divergence time for a single branch in the phylogeny (Cutter 2008) and represents the hypothetical scenario where the whole Helitron sequence has been conserved (or ‘static’) since the HT events. We considered an *μ* value of 3.0 × 10^-9^ mutations per generation per site per haploid genome for *D. rhopaloa, C. vestalis* and *B. mori*, based on the direct measures conducted on four insect species from three different orders (Keightley *et al.* 2014, 2015, Yang *et al.* 2015, Liu *et al.* 2017) and on the possibility that insects might have the same mutation rate (Liu *et al.* 2017). For alignment and estimation of evolutionary divergence between sequences (*K*) we used the software MEGA7 (Kumar *et al.* 2016). As the *T* values are given in number of generations, and we were interested in estimate the HT dates in millions of years (*mya*), two averages for the number of generations per year (gen/y) were used: one for the *D. rhopaloa - C. vestalis* (14 gen/y) and another for the *C. vestalis - B. mori* (11 gen/y) HT event. They were based on the known estimates for each taxon: ~ 10 gen/y for the *D. melanogaster* species group (Cutter 2008, McDonald and Kreitman 1991, Tochen *et al.* 2014, Asplen *et al.* 2015), ~ 18 gen/y for Braconidae (Nikam and Sathe 1983, Nikam and Pawar 1993), and ~ 4 gen/y for *B. mori* (Maekawa *et al.* 1988, Reddy *et al.* 1999).

### Data availability

The authors state that all data necessary for confirming the conclusions presented in the article are represented fully within the article.

## RESULTS AND DISCUSSION

### Segment c35 from *Cotesia vestalis* bracovirus is a Helitron autonomous TE

The segment c35 (CvBV_c35) (gb: HQ009558.1) from *Cotesia vestalis* bracovirus (CvBV) has 5,667 bp and contains a single ORF, previously described as a DNA helicase of unknown origin that has not been reported in other PDVs (Chen *et al.* 2011). Indeed, its annotated amino acid sequence (AEE09607.1) has a C-terminal Pif1-like helicase domain which belongs to the P-loop containing Nucleoside Triphosphate Hydrolases (P-loop_NTPase) superfamily. Because the Pif-like domain covers only a relatively small portion of the total amino acid sequence (< 30%) and it is absent in all known PDVs, we further investigated the identity of this protein.

We found that the N-terminal half of the protein contains features of a Rep domain which is typical of the Rep-Helicases (Rep/Hel) from Helitron TEs (Kapitonov and Jurka 2001). Subsequent analysis revealed the presence of all three conserved motifs present in the Rep catalytic core (Kapitonov and Jurka 2007), confirming the identity of this protein as a Rep/Hel (Figure 1). This group of proteins presents a basic structure consisting of a N-terminal HUH endonuclease domain (Rep) and a C-terminal helicase domain (Hel) and is used by Helitrons during their rolling-circle (RC) transposition (reviewed in Thomas and Pritham 2015).

In addition to a Rep/Hel gene, all autonomous Helitrons have conserved structural features at their termini which are also involved in the RC transposition mechanism (Thomas and Pritham 2015, Grabundzija *et al.* 2016). Thus, after we determined the Rep/Hel identity of this coding sequence, we verified if the rest of the TE was present in segment c35. By comparing CvBV_c35 and Helitron sequences retrieved from Repbase (Jurka *et al.* 2005) we could determine the structure of the whole element, with its 5’- and 3’-termini flanking the Rep/Hel ORF. This Helitron, hereafter named Hel_c35, has 5,294 bp, and covers most (~ 93.4%) of the viral segment c35 (Figure 1C). Because this element contains an intact ORF and all the structural hallmarks of Helitrons (see *Materials and Methods*), it should be classified as an autonomous TE, according to the definition proposed by Wicker *et al.* (2007). By contrast, non-autonomous Helitrons do not encode a Rep/Helicase, despite sharing common structural features with their autonomous counterparts (Thomas and Pritham 2015). Although Hel_c35 fits the definition of an autonomous TE, it is important to note that there is not enough evidence to assert whether the element is currently active. Only a small 373 bp region within segment CvBV_c35, but outside the Hel_c35 sequence, lacks significant homology with Helitrons. Moreover, we found the same 373 bp sequence (100% identity) as part of a different CvBV segment (CvBV_c19), indicating that this sequence probably has a viral origin. This 373 bp viral sequence does not display significant similarity with any known transposable elements or insect genome available to date.

The analysis performed with the CENSOR tool (Kohany *et al.* 2006) against the whole Repbase repeat library, using Hel_c35 as a query, indicated that this element has high nucleotide sequence identity (70-97%) with Helitrons from Diptera, especially Drosophila species (Figure S1). Furthermore, the Rep/Hel amino acid sequence from Hel_c35 display a similarity of ~ 73% over the whole Rep/Hel from Helitron-5_DRh (1072 aa), a different Helitron from *D. rhopaloa* (data not shown). Although Hel_c35 displays high nucleotide sequence identity with these Helitrons, they share less than 80% similarity in their last 30 bp at the 3’-terminus. Therefore Hel_c35 does not belong to any described family, according to the proposed classification of Helitrons (Thomas and Pritham 2015).

Helitron insertions in PDV genomes have been reported before, including in CvBV (Thomas *et al.* 2010, Guo *et al.* 2014, Coates 2015). It is worth mentioning that some of these studies describe Helitron insertions in the genomes of *Cotesia plutellae* and *Cotesia plutellae* bracovirus (CpBV), which are junior synonyms of *C. vestalis* and CvBV, respectively (Shaw 2003). However, these elements are non-autonomous, and display low sequence identity (< 80%) with Hel_c35, including on their terminal sequences (data not shown), thus belonging to a distinct Helitron family.

### Searching the CvBV_c35 proviral locus

PDV circles originate from integrated copies in the wasp's genome. These integrated segments are present in genomic regions known as proviral loci (Gundersen-Rindal *et al.* 2013, Strand and Burke 2013, Drezen *et al.* 2014). In order to determine the CvBV_c35 proviral locus, we Blast-searched CvBV_c35 (HQ009558.1) in the *C. vestalis* sequenced genome. There are currently two lineages of *C. vestalis* with assembled genomes: isolate ANU101 (BioSample: SAMN03273265) from South Korea and isolate 20120220 (BioSample: SAMN04378091) from China. Both strains are distinct from the one used by Chen *et al.* (2011) to obtain the viral segment CvBV_c35 (HQ009558.1). Although we found a hit with 100% similarity over ~ 40% of the query length in isolate 20120220, none of the results from this lineage contained a complete copy of CvBV_c35, and thus they were not included in the following analysis.

The best hit (gb: JZSA01007369.1) corresponds to contig 7377 from isolate ANU101 and contains a Helitron copy with ~ 99.9% nucleotide identity to the Hel_c35 element present in CvBV_c35 (Figure 2). Despite the high sequence similarity, this putative CvBV_c35 proviral sequence contains two deletions, one of 136 bp (between positions 4,780-4,915 of the Hel_c35 query sequence), and another corresponding to the whole 373 bp viral sequence within segment CvBV_c35. Thus, the similarity of this proviral copy with CvBV_c35 is limited to the Hel_c35 sequence. Curiously, Blast searches in the genomes of both wasp lineages, using this 373 bp viral sequence as a query, retrieved one hit (100% identity) in a locus that corresponds to CvBV_c19, only in the genome of isolate 20120220. Hence, this 373 bp sequence appears to be absent in the isolate ANU101, even though proviral segment CvBV_c19 (where this sequence was also found) is present in this genome.

**Figure 2.**
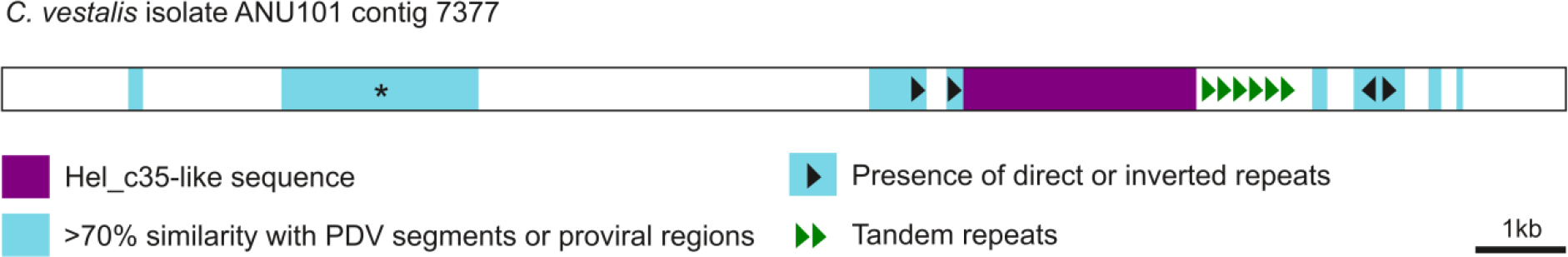
Putative CvBV_c35 proviral locus in the *C. vestalis* genome (strain ANU101). This locus contains a sequence very similar (~ 99.9% identity) to Hel_c35. The region is marked by the presence of several sequences similar to other polydnaviruses (in blue), and the presence of direct and inverted repeats (black arrowheads). The asterisk denotes a PDV conserved region with a complex array of direct, inverted and tandem repeats.

Instead of the 373 bp viral sequence flanking its termini, this Hel_c35 copy in the *C. vestalis* genome presents a 396 bp sequence which is tandemly repeated 6 times after the Helitron 3’-end (Figure 2). These repeats are not abundant in the *C. vestalis* genome, being present as less than 100 copies distributed over ~ 20 short arrays containing on average four repeats. We did not find these repeats in other hymenopteran genomes, including *Microplitis demolitor*, which belongs to the same subfamily as *C. vestalis* (Microgastrinae), and currently is the closest wasp species with a sequenced genome. Moreover, we noticed that upstream the Helitron 5’-terminus there are two long direct repeated sequences with 376 bp each and ~ 99.5% identity, separated by 426 bp, and with similarity to BV sequences (Figure 2 and Table S1). Those differences prompted us to analyze the flanking regions of this Hel_c35 copy up to several kbp upstream and downstream, to identify other possible PDV-related sequences.

We found eight motifs that are conserved in many PDV-related sequences, including several BV circles and proviral regions flanking segments (Figure 2 and Table S1). Five of the eight conserved motifs are also found within or nearby nudiviral-like genes from *Cotesia congregata* bracovirus (CcBV). Despite not being encapsidated, these genes are involved in BV replication and production of structural viral proteins (Strand and Burke 2013, Drezen *et al.* 2014). The fact that some of these motifs are found in both segment and nudiviral loci, and others are exclusive to segment loci, might be related to their common or specific functions in BVs, respectively. For example, common motifs could have a role in proviral replication, and specific motifs in circle excision, encapsidation and transcription. All the conserved sequences analyzed here have an average AT content of 71% (62.7-80%), and contain internal palindromic regions that could form hairpin folds with variable sizes. Additionally, two of these motifs around the Hel_c35, one upstream and the other downstream, contain long inverse repeats that could also form stem-loop structures (Figure S2). Similar features have been described in flanking and intermediate sequences of other BV proviral loci, and are thought to be involved in viral replication (Louis *et al.* 2013, Burke *et al.* 2015).

The presence of several motifs flanking this Hel_c35 in the *C. vestalis* genome suggests that this copy was probably inserted on a proviral locus. If true, why this Hel_c35 copy is different from the one present in the viral genome? This inconsistency might be explained if this Hel_c35 insertion does not correspond to the main proviral CvBV_c35, but to a paralogous segment that could be either active (it only produces a small part of c35 circles) or inactive (such as a ‘pseudo-segment’). A large portion of PDV genes found in viral segments belong to multigenic families, and segment reintegration, locus duplication/deletion, and gene gain/loss, are common features during PDV evolution (Herniou *et al.* 2013, Burke and Strand 2012a).

Another possibility, which perhaps is more plausible, is that CvBV_c35 is polymorphic in the different lineages used for the genome sequencing of *C. vestalis* (BioSample: SAMN03273265) and CvBV (HQ009558.1). In addition to different CvBV_c35 segments in distinct lineages, this polymorphism could also include lineages with no CvBV_c35 encapsidated circles. Before the work conducted by Chen *et al.* (2011), the CvBV (referred as CpBV) genome was partially sequenced from a wasp strain of South Korea (Choi *et al.* 2009), and no circles containing a Rep/Hel gene were detected. Although this is a partial genome sequence, the missing CvBV_c35 could be a consequence of its real absence in the CvBV particles from this wasp strain. As we described above, the best candidate for a proviral CvBV_c35 sequence also belongs to a wasp lineage from South Korea, in contrast to the CvBV_c35 viral sequence (HQ009558.1), which is derived from a wasp strain of China. Hence, the differences between the wasp and BV genomes might reflect a polymorphic CvBV_c35 which, depending on the lineage, exists as a proviral segment that generates encapsidated viral circles, or as an ‘ancestral’ form with the Hel_c35 in a proviral region, but not able to produce viral particles. Although the confirmation of this hypothesis of polymorphism would require the genome sequencing of different *C. vestalis* strains and their respective BVs, it is worth mentioning that there is evidence of significant variation between BVs from different wasp populations of the same species (Rincon *et al.* 2006, Branca *et al.* 2011).

### Rep/Hel transcripts in a lepidopteran host parasitized by *Cotesia chilonis*

The data described here so far indicate a Helitron-BV fusion. In this context, it is important to investigate when, during the evolution of BV, this event happened. Apart from CvBV, we did not find the Hel_c35 Rep/Hel ORF in any other PDV sequenced genome available to date, including those from two other bracoviruses present in two *Cotesia* species: *Cotesia congregata* bracovirus (CcBV) and *Cotesia sesamiae* bracovirus (CsBV). Although there are no other *Cotesia* sequenced genomes available, Wu *et al.* (2013) described the influence of *Cotesia chilonis* on its lepidopteran host *Chilo suppressalis* during parasitization, by analyzing the transcriptome of fat body and hemocytes in the larvae of this moth. In addition to host genes, they identified 19 unique sequences associated with PDVs, which were classified as *Cotesia chilonis* bracovirus (CchBV) transcripts; so we further searched for sequences similar to Hel_c35 on their Short Read Archive (SRA) (accession: SRR651040) and Transcriptome Shotgun Assembly (TSA) (gb: GAJS00000000) databases in order to pinpoint the approximate phylogenetic position of the Hel_c35 integration into the ancestral BV genome.

We detected ~ 70 reads with > 80% similarity, and ~ 50 reads with > 90% similarity with the Hel_c35 ORF in the SRA, covering different regions of its sequence (Figure 3). These reads were mainly located around two Hel_c35 ORF regions extending 223 and 608 bp, which were assembled as Unigene42046 (gb: GAJS01040222.1), and Unigene57509 (gb: GAJS01055664.1), respectively (Figure 3). Interestingly, searches on the genomes of *C. suppressalis* and *Amyelois transitella* (both from the Pyraloidea superfamily) using Unigene42046 as a query retrieved sequences with only 74-85% identity. On the other hand, the same Unigene has 99% sequence identity with the Rep/Hel ORF from CvBV_c35 (Figure S3). The high similarity of this transcript with the CvBV segment, and its likely absence from the genome of *C. suppressalis*, suggests that this transcript is derived from CchBV.

**Figure 3.**
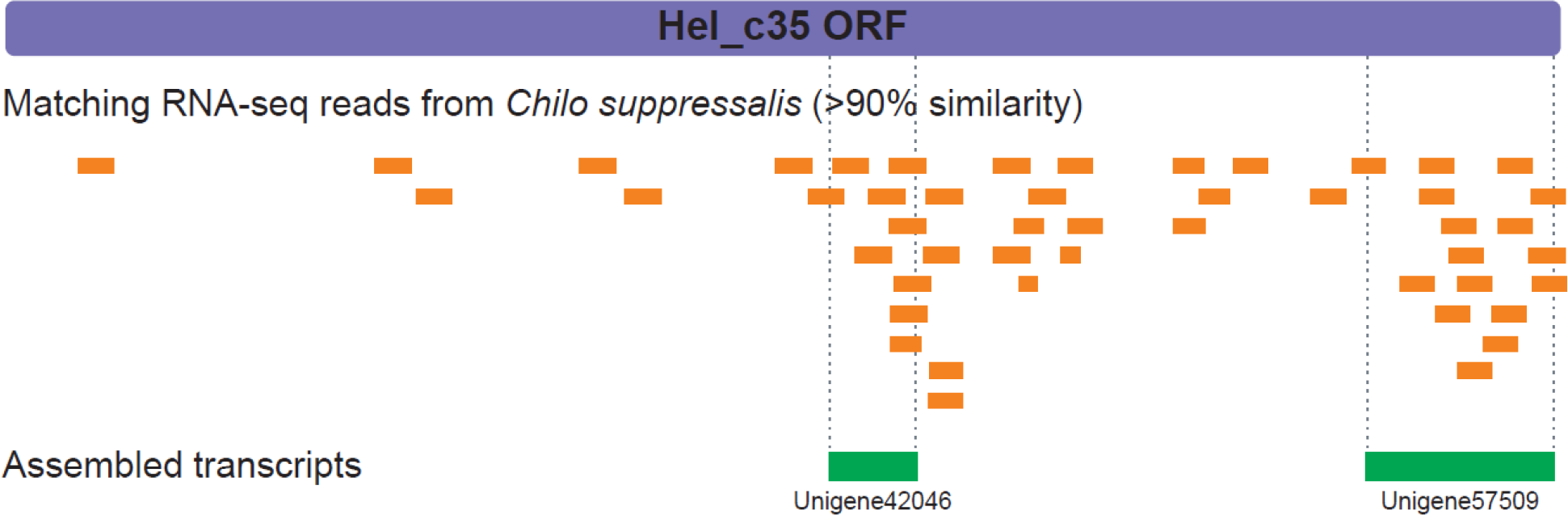
Transcripts from a lepidopteran host parasitized by *Cotesia chilonis*. Reads and assembled Unigenes from *Chilo suppressalis* parasitized by *Cotesia chilonis*, and displaying sequence similarity (> 90%) with Hel_c35 regions.

We then applied the information about the Rep/Hel distribution into the *Cotesia* phylogeny reconstructed by Michel-Salzat and Whitfield (2004). Considering the topology of the species represented in Figure 4, we might assume one of the two following scenarios. The first is that a homologous bracovirus segment with a Rep/Hel gene was independently lost twice during *Cotesia* evolution, one in the *C. congregata* and the other in the *C. sesamiae* clade (Michel-Salzat and Whitfield 2004). The second is that two independent Helitron integration events occurred, one in CvBV and the other in CchBV. The hypothesis of two integration events involving two different Helitron families would explain the disparity of the Hel_c35 ORF sequence similarity with Unigene42046 (99%) and Unigene57509 (71%). In this case, both Unigenes would be part of one single ORF from a Helitron family that is distinct from Hel_c35. This hypothesis is also supported by our date estimate of when Hel_c35 was first inserted in the *C. vestalis* genome (< 1 mya) (see topic ‘*Tracing an evolutionary pathway of Hel_c35 in insect genomes*’), which appears to be too recent to have happened before the divergence of the two species. Nevertheless, the genome sequencing of *C. chilonis* and CchBV will be necessary to test (i) the identity between Hel_c35 and Unigene42046, (ii) the presence of this TE within CchBV, and (iii) the homology of this insertion with the one found in segment CvBV_c35.

**Figure 4.**
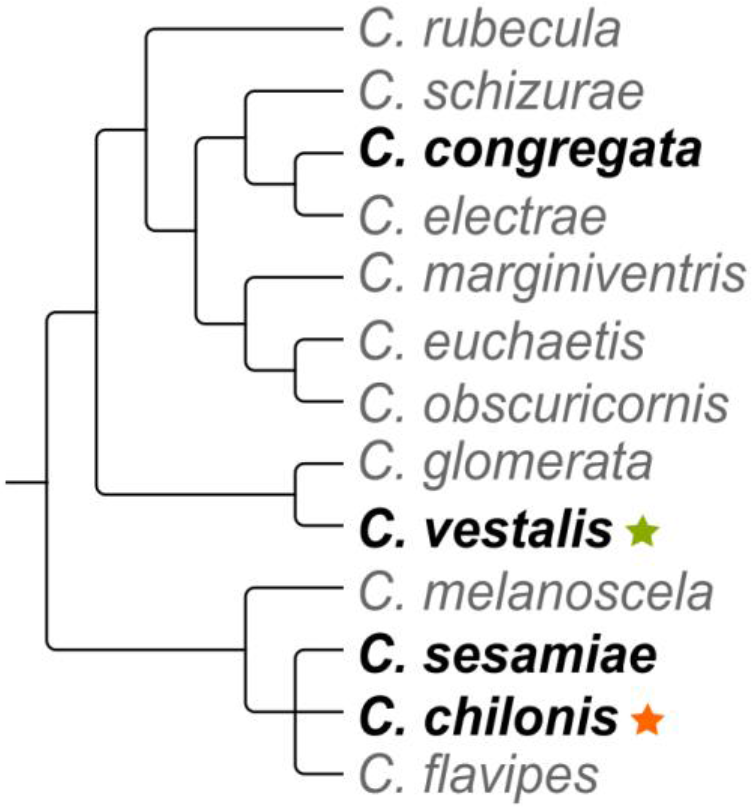
*Cotesia* phylogeny. The phylogenetic position of four *Cotesia* species (in bold) from which we investigated the presence of Hel_c35 in the respective BV genomes. Presence of Hel_c35 in *C. vestalis*, and putative presence of Hel_c35 in *C. chilonis* represented by green and orange stars, respectively. Adapted from (Michel-Salzat and Whitfield 2004).

### Horizontal transfer of Hel_c35 between three insect orders

To better characterize the evolution of the Helitron Hel_c35, we searched for sequences displaying high similarity with the ORF's Rep domain from CvBV_c35 (File S1) in all arthropod sequenced genomes available to date. A nucleotide alignment containing all obtained sequences is shown in Figure S4. We then used these sequences to construct a phylogenetic tree (Figure 5, see *Materials and Methods*).

The resultant tree revealed some strikingly incongruences with the species' phylogeny (Figure 5). Most notably, Hel_c35 from *Bombyx mori, Drosophila rhopaloa* and *Drosophila ficusphila* were allocated in the same clade as the Hel_c35 from *Cotesia vestalis* bracovirus (CvBV). In fact, the nucleotide identity between the Hel_c35 sequence from CvBV, and the Hel_c35 from *B. mori, D. rhopaloa* and *D. ficusphila* are ~ 99.7%, ~ 96% and ~ 94%, respectively. It is important to mention that we did not find Hel_c35 in any other hymenopteran species, even though more than forty genomes from this insect order were used in our search. Because the marked phylogenetic incongruences and the extreme discontinuous or “patchy” distribution of Hel_c35 are indicative of horizontal transfer (HT) events (Silva *et al.* 2004, Wallau *et al.* 2012), we decided to further investigate this possibility.

For a HT event to be inferred, the species distributions also should be considered, as the candidate taxa must overlap geographically at some degree in order to a HT event to occur (Loreto *et al.* 2008, Carareto 2011). Thus, we classified each taxon in the phylogeny to one of eight major geographical regions, based on their known distributions. The addition of this information revealed interesting aspects that may help to explain some of the main inconsistencies found in the phylogeny (Figure 5). First, among the ten taxa closest to the Hel_c35 query sequence, seven are native from eastern/southestern Asia, including the three species that group immediately with the Hel_c35 from CvBV (*B. mori, D. rhopaloa* and *D. ficusphila*). Second, the positioning of *Bactrocera dorsalis* (Tephritidae) within a well-supported clade of Drosophila species, instead of a position closer to other tephritid fruit flies, also coincides with their common distribution in southeastern Asia. Third, the incongruent grouping of *Calycopis cecrops* (Lepidoptera) with *Drosophila willistoni* (Diptera), involve two species with geographic overlapping in southeastern United States. Fourth, even though *Drosophila mojavensis* and the lepidopteran *Amyelois transitella* belong to different insect orders, they are grouped together in a well-supported clade and overlap geographically in southwestern United States. Thus, the geographical distribution of the analyzed taxa supports the HT hypothesis as an explanation for at least some of the phylogenetic incongruences, especially the one directly related to CvBV. Other incongruences outside the immediate CvBV clade might also represent bona fide HTs; however, each of them requires a careful analysis that is beyond the scope of the present work.

For HT events, the spatial overlapping of candidate species must be associated with at least some degree of ecological overlapping (Loreto *et al.* 2008, Carareto 2011). Accordingly, braconid wasps interact very closely with several insect host orders, including lepidopteran and Drosophila species (Quicke 2015, Wharton 1993) and as a rule, these connections reach the cellular and even the chromosomal level during wasp oviposition and bracovirus infection of host cells (Gundersen-Rindal *et al.* 2013). It is noteworthy that these extremely direct interactions are not occasional, but a fundamental part of the braconid wasps' life cycle.

**Figure 5.**
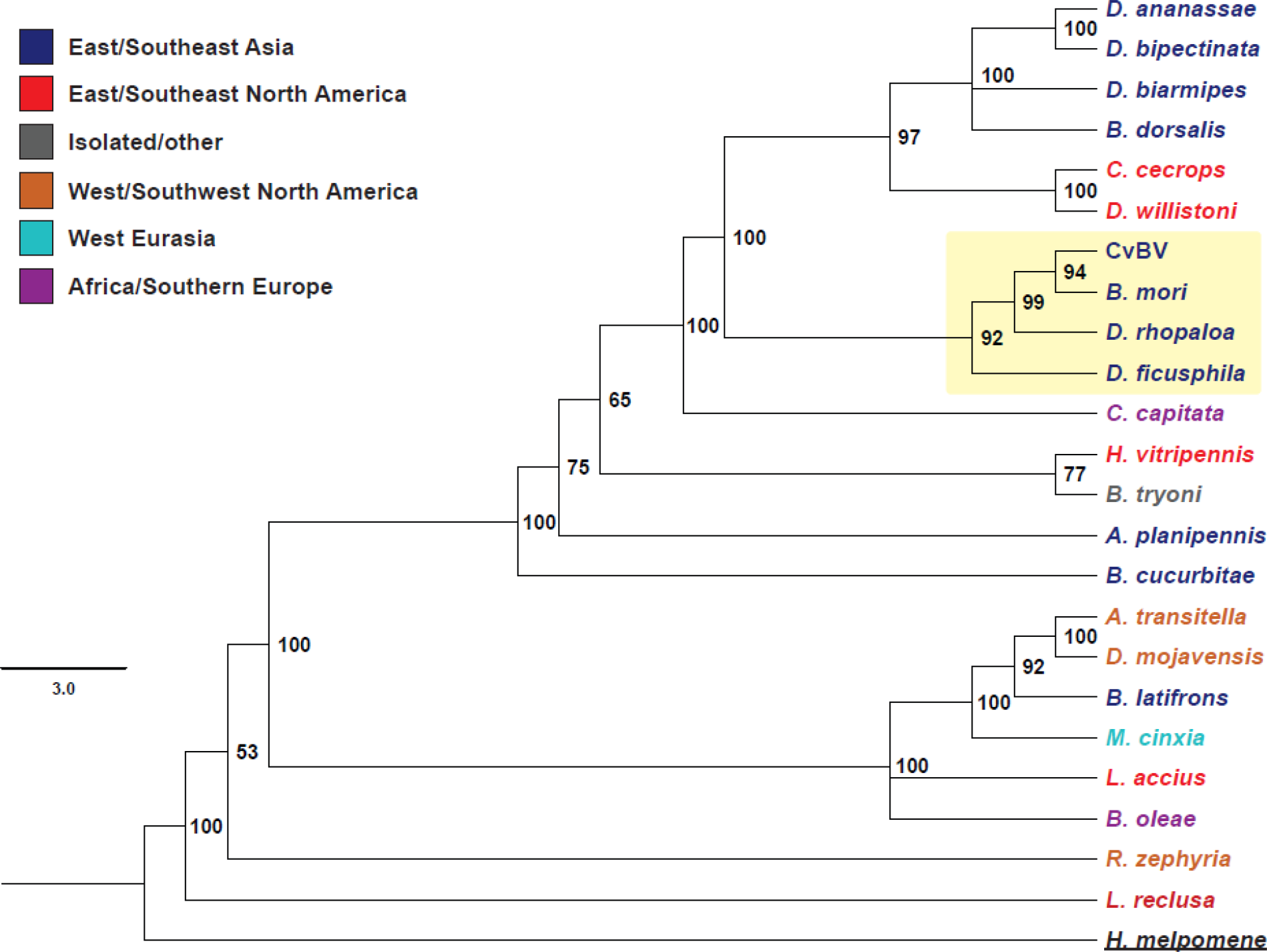
Phylogeny of Hel_c35 core Rep domain sequences in several insect genomes. The phylogeny was built using Bayesian inference (see *Materials and Methods*) on a aligned Rep region with ~ 838 bp (Figure S4). Species name colors indicate the major geographical regions in which these species are found. CvBV clade is highlighted in yellow. A list of species with their orders and accession numbers is given in Table S2.

In summary, our results suggest at least two HT events involving Hel_c35: one between a Drosophila species and *C. vestalis* and another between *C. vestalis* and *B. mori*, which probably occurred in this respective order, as indicated by the phylogeny. This hypothesis is supported by four lines of evidence: (i) the marked incongruence between host and TE phylogeny, (ii) the patchy distribution of this TE on the main taxa involved, (iii) the high sequence identity between Hel_c35 copies from different insect orders, and (iv) the spatial/ecological overlap among the candidate species.

### Tracing an evolutionary pathway of Hel_c35 in insect genomes

Because our constructed phylogeny with Hel_c35 sequences suggests HT events between *D. rhopaloa, C. vestalis* and *B. mori*, we estimated when these events occurred using the equation for divergence time (*T* = *K*/2*r*) on non-coding sequences (~ 750 bp) from Hel_c35 copies of these species. We also set minimum and maximum date thresholds, supposing different evolutionary constrains acting on these sequences (see *Materials and Methods*). This analysis can also help to infer the order of the two HT events which, according to the phylogeny, appear to have occurred from *D. rhopaloa* to *C. vestalis* (represented by CvBV), and then from *C. vestalis* to *B. mori*. The results give an approximate date of 0.862 mya (0.574 – 1.15 mya) for the *D. rhopaloa – C. vestalis* HT, and 0.211 mya (0.141 – 0.282 mya) for the *C. vestalis – B. mori* HT. These values are striking, considering that *C. vestalis* (Hymenoptera) diverged from *D. rhopaloa* (Diptera) and *B. mori* (Lepidoptera) approximately 325 mya and these last two species diverged from each other approximately 272 mya (Kumar et al. 2017).

Based on our results, we suggest the following scenario to explain the observed distribution of Hel_c35 copies among insect genomes. The transposable element Hel_c35 belongs to an undescribed Helitron family from southeastern Asian Drosophila species and, more specifically, from the oriental subgroup cluster of the *Drosophila melanogaster* species group. This taxon comprises the largest number of species with Hel_c35-related sequences clustering with CvBV, and displays a topology roughly coherent with the group's phylogeny (Seetharam and Stuart 2013). Although Helitrons constitute a large portion of Drosophila genomes across different groups (Yang and Barbash 2008, Dias et al. 2015, de Lima et al. 2017), the distribution of element Hel_c35 appears to be originally restricted to a specific clade within the oriental subgroup cluster. Initially, a Hel_c35 element from a Drosophila species in the *rhopaloa* subgroup was horizontally transferred into the ancestral genome of *C. vestalis* ~ 0.862 mya, probably during parasitization. After genome integration, Hel_c35 transposed into the CvBV proviral genome in a locus responsible for segment production, also becoming part of the encapsidated genome of this bracovirus. Then, a second HT event occurred ~ 0.211 mya, this time involving the transfer of Hel_c35 from *C. vestalis* to *B. mori*, presumably through CvBV_c35 circles (Figure 6). If after entering the proviral locus, Hel_c35 ‘fused’ gradually with CvBV, the observed sequence inconsistencies between CvBV_c35 from different *C. vestalis* lineages could represent incomplete stages of this fusion on different wasp strains. In that case, the formation of segment CvBV_c35 would have been completed after geographical divergence of the lineages used for wasp (S. Korea) and BV (China) genome sequencing.

**Figure 6.**
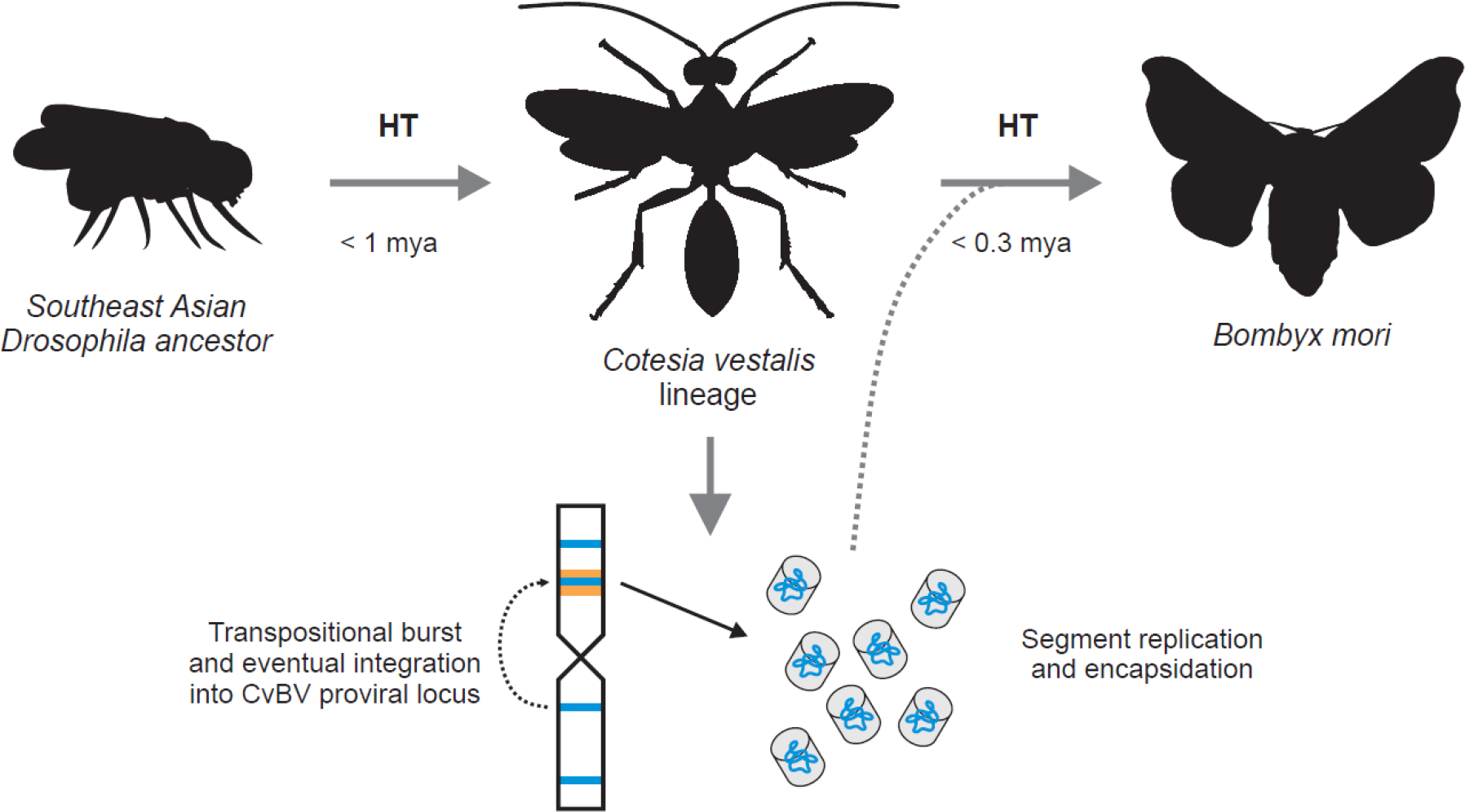
Hypothetical evolutionary history of Hel_c35. This TE was already present in the ancestor of the oriental subgroup of the *melanogaster* species group. Less than one million years ago (mya), possibly during non-specific parasitization of *Drosophila rhopaloa*, Hel_c35 was horizontally transferred to the *Cotesia vestalis* lineage. Following transpositional activity in *C. vestalis* chromosomes, Hel_c35 eventually inserted into an active proviral locus and begun being replicated and encapsidated as the segment c35. Now replicated to high copy numbers and able to infect cells, it was horizontally transferred again (< 0.3 mya) to *Bombyx mori*, a likely Lepidopteran host of *Cotesia vestalis*.

To hold true, the above hypothesis might require at least two assumptions that should be addressed. First, that *C. vestalis* is or at least was until recently capable to parasitize Drosophila species, in addition to their typical lepidopteran hosts. This hypothesis may seem improbable at a first sight, because all hosts of Microgastrinae wasps described to date are Lepidoptera larvae (Quicke 2015). Indeed, the major radiation dates for both taxa seem to coincide, reinforcing the idea that Microgastrinae have evolved as a group of specialized Lepidoptera parasitoids (Banks and Whitfield 2006). However, it is also important to note the remarkable host diversity and variability within the Braconidae family. For instance, the subfamilies Alysiinae and Opiinae are endoparasitoids of many cyclorrhaphous Diptera, including Drosophila and tephritid fruit flies (Wharton 1993, Carton *et al.* 1986), and several wasp species in the Braconinae subfamily can attack both dipteran and lepidopteran hosts, depending on the location and time of the year (Žikić *et al.* 2012, Gadallah and Ghahari 2015). The Exothecinae subfamily contains parasitoids of several insect orders: for instance, *Colastes braconius* can parasitize dipteran, lepidopteran, coleopteran and even hymenopteran species (Shaw and Huddleston 1991). Other ecological aspects are also relevant if we consider the host range and specificity within the *Cotesia* genus. The first is that, even though *Plutella xylostella* is commonly described as the main host of *C. vestalis* in the literature, this wasp is capable to parasitize a wide range of Lepidoptera families and superfamilies, which are extremely diverse in their ecology and morphology (Cameron and Walker 1997, Malysh *et al.* 2016).

It is also worth mentioning that, although to our knowledge there is no report of *C. vestalis* parasitization on *B. mori*, the close related species *Cotesia glomerata* can use this moth as a host (Sathe and Jadhav 2001), and *Cotesia dictyoplocae*, which is in the same species group of these two wasps, parasitizes moths of the Bombycoidea superfamily (Gupta *et al.* 2016). Additionally, the present work is not the first to report a HT between *C. vestalis* and *B. mori* (e.g. Coates 2015, Zhang *et al.* 2016a, Zhang *et al.* 2016b), indicating that *C. vestalis* could indeed attack this lepidopteran species, and might also parasitize a wider variety of hosts.

Secondly, the known hosts of a parasitoid wasp do not necessarily correspond to their actual host range, and may simply represent the commonly attacked species of which successful parasitization is more likely to ensue. It has been suggested that unusual conditions could induce attacks outside the suitable host range, and even result in successful parasitization of unsuitable hosts, notwithstanding their rarity (Heimpel *et al.* 2003, Quicke 2015). It is also noteworthy that some Drosophila species phylogenetically close to *D. rhopaloa* can be resistant to attacks from the wasp *Asobara japonica*, probably because of a long-continued interaction with braconid parasitoids (Ideo *et al.* 2008, Furihata *et al.* 2016). Additionally, on southeast Asia, two Drosophila species closely related to *D. rhopaloa* breed on plants that are also commonly used as food by Lepidoptera larvae (Suwito *et al.* 2002), which could facilitate an encounter between *Cotesia* wasps and unusual dipteran hosts. Hence, it is not difficult to conceive that *Cotesia* wasps attack, and sometimes can successfully parasitize Drosophila larvae, even if these interactions may be rare.

Finally, it is important to note that any current HT analysis involving distant related taxa is subject to different interpretations in the future. That is because the available genome sequences only represent a small sample of the whole taxa diversity with the potential to be part of the described events. Thus, with an increasing number of species with sequenced genomes, there will always be the possibility of more accurate and complete descriptions for HT events.

### PDV-Helitron exaptation?

There are several reports of TE integrations in PDV (e.g. Drezen *et al.* 2006, Dupuy *et al.* 2011). Moreover, Helitrons have already been found in the genomes of *Cotesia* species and in their respective PDVs, and were also involved in HT events (Thomas *et al.* 2010, Guo *et al.* 2014, Coates 2015). However, all these insertions represent non-autonomous short or fragmented elements. To our knowledge, we describe the first instance of a putative autonomous Helitron that not only integrated the PDV genome, but effectively became a viral segment.

Exaptation refers to features originally evolved for some function but that were later co-opted for a different role (Gould and Vrba 1982). Repetitive DNAs like TEs can also take part on this process (Brosius and Gould 1992), as revealed by many studies in eukaryotes (recently reviewed by Chuong *et al.* 2017). Although there is evidence for the role of TEs on PDV evolution, including the exaptation of a gene apparently derived from from a retroelement (reviewed in Burke and Strand 2012b), we report the first evidence of an intact TE that appears to have been co-opted by a bracovirus. The main reasons for this suggestion are: (i) this element has all the features of an autonomous TE; (ii) it has conserved an intact structure in the genome of *C. vestalis* for the last ~ 1 my, despite the low number of copies; (iii) the element occupies almost 94% of segment CvBV_c35, which is one of the most replicated circles of this virus (Chen *et al.* 2011); (iv) it contains the only ORF in CvBV_c35, and all known PDV segments have coding sequences, with a few exceptions (Burke *et al.* 2014); (v) the CvBV_c35 portion outside Hel_c35 has only 373 bp, and does not contain any ORF or conserved sequences found in other PDVs (apart from segment CvBV_c19). Although it is possible that this short 373 bp sequence has a functional role (e.g. as an encapsidation signal), the whole PDV segment CvBV_c35 is essentially a Helitron. Furthermore, we found a single full copy of Hel_c35 in the *C. vestalis* genome, even though *D. rhopaloa* and *B. mori* contain several highly similar partial copies (data not shown). This unique arrangement for a TE in a eukaryote genome further suggests that segment CvBV_c35 has been kept as an intact Helitron by selective constraints.

Supposing that Hel_c35 is not just a selfish element within the CvBV genome, but also play an active role in this bracovirus, an import question emerges: what could be the advantage of having a Helitron in a PDV genome? We have three non-mutually excluding hypotheses to explain this question.

The first hypothesis is that this Helitron Rep/Hel could have been exaptated by CvBV for its helicase function, to aid the amplification of viral replication units or circles in wasp calyx cells. Most viral genes thought to be essential for BV replication have not being identified, with a few exceptions, like a nudiviral helicase and a fen-like flap endonuclease (Herniou *et al.* 2013). The Pif1 family of helicases (which include the Hel domain in Rep/Hel proteins) and the flap endonucleases are important to process DNA secondary structures like hairpin or fold-back substrates during replication (Pike *et al.* 2010, Balakrishnan and Bambara 2013). As we have mentioned, TA-rich inverted repeats and other palindromic motifs capable of forming hairpins are found throughout the proviral genomes of BVs, and probably serve as replication origins (Louis *et al.* 2013, Burke *et al.* 2015). Although possible, this hypothesis does not explain why the Rep/Hel ORF is part of the CvBV encapsidated genome, as that would not be necessary for segment replication in calyx cells.

The second hypothesis is that the whole Helitron (with its non-coding sequences) is used for specific segment CvBV_c35 amplification in calyx cells. That could explain why CvBV_c35 constitutes one of the most abundant circles in CvBV, despite being recently incorporated in the viral genome. This ‘auto-replication’ would be useful if PDV capsids also play a role on virulence, independently of the genes they carry. In that case, abundant segments could be selected for their ability to assemble more capsids that are later injected into the host. This hypothesis is based on the observation that *Microplitis demolitor* produces some BV encapsidated segments with no apparent coding sequences (Burke *et al.* 2014), and several parasitoid wasp species, including braconids, use virus-like particles (VLP) with no detected PDV sequences as virulence factors against their hosts (Herniou *et al.* 2013, Furihata *et al.* 2016, Rizki and Rizki 1990). In contrast to the first hypothesis, this scenario also predicts the conservation of Hel_c35 non-coding sequences, as they would be at least partially necessary for segment CvBV_c35 to take advantage of the Helitron rolling-circle replication (RCR) mechanism. That is because Rep/Hel proteins recognize Helitron terminal sequence motifs to start and finish RCR correctly (Grabundzija *et al.* 2016). Although the sequences used in our analysis display slightly higher nucleotide conservation on coding regions in comparison with non-coding regions (data not shown), the exact level of sequence stability necessary for Rep/Hel recognition and RCR viability of Helitrons is not well understood.

The third hypothesis is that the Rep/Hel gene from CvBV_c35 can be used to replicate no only its own segment, but also other CvBV circles, once within host cells. Like the second hypothesis, this explanation considers that the Rep/Hel is part of the BV encapsidated genome, but in this case, the Rep/Hel protein would also amplify other CvBV circles after their injection in the host. This mechanism of circle replication in host cells could further increase the number of CvBV genes, and therefore enhance virulence. This scenario would be particularly interesting to investigate experimentally in the future, as all PDVs analyzed to date only express replication genes in the calyx cells of wasps, and apparently do not replicate after encapsidation and injection in host cells (Gundersen-Rindal *et al.* 2013, Bézier *et al.* 2009). Thus, the hypothesis of Hel_c35 exaptation by CvBV for a replication-related function is plausible, even though the replication mechanism of bracoviruses is not fully understood.

### Concluding remarks

There are several reports of TE integrations in PDVs. However, all these cases involve classical TE insertions within PDV genomes, mostly of non-autonomous and fragmented elements. In contrast, our study revealed an autonomous Helitron within a bracovirus segment, which is not only an integrated copy, but a TE covering almost entirely a viral circle. The segment CvBV_c35 is effectively an intact Helitron, and our results suggest that this TE might have been recently co-opted by CvBV, probably for its helicase and/or rolling-circle replication function. This PDV-Helitron fusion points to a new type of relationship between TEs and viruses, which should be further investigated. In addition, our data reinforces the idea of PDVs as effective agents of horizontal transfer (HT), and of Helitrons as one of the TEs most commonly involved in those events. Specifically, we reported two consecutive transfers of Hel_c35 across three insect orders.

The number of reported HT events in eukaryotes has been growing recently, although the probability of their occurrence must be very low, as they require the presence of several biological features and conditions. Notably, the tripartite parasitoid system composed by braconid wasps, PDVs and hosts fulfill most of those requirements (Venner *et al.* 2017), being exceptional candidates for the study of HT in eukaryotes. Indeed, there are several reports of HT events involving insect species associated with these parasitoid systems (Drezen *et al.* 2017). As pointed out by Quicke (2015), because the number of estimated Microgastrinae species ranges between 16,000-40,000 and apparently, all of them have associated PDVs, this could be the biggest known group of viruses. The future genome sequencing of more PDVs and wasp species will help to understand the real importance of HT for the evolution of parasitoids and their hosts. Furthermore, the investigation of events like this can contribute to ecological analyses, by revealing potential hosts and cryptic interactions not detected in field studies.

## ACKNOWLEDGEMENTS

This work was supported by grants from “Fundação de Amparo à Pesquisa do Estado de Minas Gerais” (FAPEMIG) (grant number APQ-01563-14), and from master and doctoral fellowships from “Coordenação de Aperfeiçoamento de Pessoal de Nível Superior” (CAPES) to PH and GBD, respectively.

